# Differential T cell receptor clonotype abundance analysis with similarity-based counts weighting

**DOI:** 10.1101/2025.03.28.645951

**Authors:** Franka Buytenhuijs, Thomas C. Smits, Ankur Ankan, Johannes Textor

## Abstract

To better understand immune responses, comparing the abundance of T cell receptors (TCRs) between conditions can provide insights into which T cells have proliferated or were involved in immune activation. This requires methods that can accurately identify significant differences in TCR-seq data. For conventional RNA-seq data, well-established differential gene expression (DGE) analysis tools such as DESeq2 and edgeR have been developed. However, applying these methods to TCR sequencing (TCR-seq) data presents additional challenges. TCR-seq data is highly sparse, overdispersed, and contains many artificial zeros, which can lead to inflated false-positive rates when using traditional approaches. While non-parametric methods like the Wilcoxon test better control false positives, they may suffer from lower statistical power.

To address these issues, we propose a novel pre-processing step for TCR-seq data using network-based local weighting based on TCR sequence similarity. This pre-processing step improves the sensitivity and reduces the false-positive rates of methods like DESeq2 and edgeR while enhancing the power of the Wilcoxon test. Through empirical analysis of both simulated and real datasets, we show that combining our pre-processing step with the Wilcoxon test achieves robust performance, outperforming traditional RNA-seq methods. This simple but powerful approach to TCR-seq analysis could help advance our understanding of adaptive immune responses.

## Introduction

Bulk and single-cell RNA sequencing allow researchers to measure and compare transcriptomes in different conditions.^1^ One of the most important and fundamental methods for analyzing RNA-seq data is DGE analysis. Broadly defined, DGE analysis refers to statistical methods that identify genes exhibiting significant differences in expression levels between groups or conditions.^2, 3^ By comparing gene expression levels between different states–such as healthy versus diseased tissues or untreated versus treated samples–DGE analysis can provide insights into the biological mechanisms that drive the observed changes. This helps researchers to understand the molecular basis of diseases, develop targeted therapies, and enhance our knowledge of fundamental biological processes.

A common strategy in DGE analysis is to fit the distribution of gene expression values to (possibly normalized) read counts. Statistical tests, such as linear regression^4^ or likelihood ratio tests,^5^are then applied to determine if the observed differences are significant. Additionally, since DGE analysis on high-throughput sequencing data involves testing thousands of observations, multiple testing corrections are necessary to reduce false positives.^6^ More advanced parametric techniques such as edgeR^7^ and DESeq2^8^incorporate negative binomial distribution models to account for the discrete and overdispersed nature of RNA-seq data. While such parametric assumptions are required to work with small sample sizes, they may lead to elevated false positive rates (FPRs) if not met.^9^ To address this issue, the rank-based Wilcoxon test has been suggested for larger sample sizes, as it is more robust in controlling the FPR and outliers while reaching comparable power to parametric approaches for medium to high sample sizes.^9^

In T cell receptor (TCR) repertoires, similar analysis is used to examine the differences in TCRs under various conditions. TCRs are found on the surface of T cells and are central to the adaptive immune system’s ability to recognize and eliminate pathogens, infected cells, and cancerous cells. Investigating differential patterns in TCR sequencing (TCR-seq) data extends DGE analysis into the immune landscape, allowing researchers to study which TCRs are upregulated in a condition, which can indicate their involvement in immune responses to certain pathogens or diseases. Rather than genes, differential analysis in TCR data focusses on differential abundance (DA) in clonality.

There is no standardized workflow for DA in TCR-seq data, but different DGE analysis tools have been applied, including edgeR^1011^,^12^ DESeq,^13^ and Wilcoxon,^14^ which is also applied by default in Immunarch.^15^ Other approaches such as Bayesian modelling have also been applied.^16^

However, DA analysis on TCR-seq data leads to additional challenges compared to DGE analysis on conventional RNA seq data. TCR sequences are highly variable,^17, 18^ and read counts can be influenced by sampling biases and differences in sequencing depth,^19^ leading to an abundance of artificial zeros in the count matrix. Importantly, these zeros do not all carry the same meaning: they can represent biological zeros (indicating absence due to biological reasons), technical zeros (caused by limitations in sequencing or processing), or sampling zeros (arising from insufficient sampling).^20, 21^ Distinguishing among these types of zeros is important for downstream analysis using DA techniques and interpretation of the results. In addition, low counts do not necessarily imply irrelevance; even lowly expressed T cell receptors, including those involved in immune responses, can be functionally important.^22–24^ Finally, like all RNA-seq data, TCR transcript counts are typically overdispersed – the variance of the counts increases with the expression level. Consequently, the presence of overdispersion complicates DA analysis, making it difficult for DA approaches to detect significant changes in the expression.

Recognizing these challenges when applying DA analysis to TCR-seq data, we propose a novel pre-processing step to adjust the TCR expression levels by normalizing read counts based on those of neighbouring sequences. It is known that T cells with receptor sequences similar to those of known antigen-binding T cells are often likely to bind the same epitopes.^25–28^ The binding of TCRs is influenced by structural and sequence motifs, meaning that slight variations in the receptor sequences can preserve the capacity to recognize a particular antigen. Consequently, clusters of TCRs with similar sequences frequently occur after an immune response, reflecting the expansion of antigen-specific T cell populations.

Our method utilizes this information by constructing a TCR sequence network, connecting sequences based on similarity, and modifying their count data using a kernel-based approach. This pre-processing addresses issues such as sampling zeros and low expression, enabling more robust detection of antigen-reactive TCRs. As this is a pre-processing step, our approach is compatible with existing DA analysis methods, providing a flexible solution for TCR-seq data. Through empirical analyses, we demonstrate that this pre-processing significantly enhances the detection of relevant signals.

## Methods

To address challenges in DA analysis of TCR-seq data, we propose a pre-processing step incorporating sequence similarity to improve the detection of low-expressing TCRs. We create similarity matrices based on different sequence comparison techniques and use them to adjust the count data before conducting DA analysis. This approach is applied to both simulated and real datasets to evaluate its effectiveness in improving the sensitivity and accuracy of TCR expression analysis.

### Data

#### Simulated data

To evaluate the performance of local weighting methods in DA analysis, we used simulated TCR-seq datasets generated with immuneSIM, a tool designed for creating T cell and B cell repertoire data.^29^ A total of 90 samples were simulated, with 45 assigned to the treatment set and 45 to the non-treatment set. Then, we incorporated additional CDR3β sequences derived from the VDJdb motif database^30^ to model the antigen-reactive sequences. Two sequence motifs, containing 80 and 57 sequences known to be reactive to Influenza A, were selected to model the reactive repertoire. Reactive sequences were sampled from the amino acid probability distribution of these motifs and replaced a random subset of the original sequences in the simulated datasets. The treatment group was designed to have a stronger immune response, modeled by a higher number of reactive sequences. In the treatment set, 206 reactive sequences were added (1.5 times the number of sequences in each motif), while in the non-treatment set, 69 sequences were added (0.5 times the number of sequences in each motif).

To simulate the proliferation of reactive T cells, we used a Galton-Watson branching process to model sequence expansion in the treatment set.^31^ In this stochastic model, each reactive sequence had a 30% probability of dividing into two daughter cells per generation step. All generated daughter cells could proliferate with the same probability in a consequent generation step. This process continued until no new offspring were generated during a generation step, signaling the end of proliferation. Sequence counts were then updated by multiplying the original values by the total number of offspring produced.

#### T cell receptor data

We applied our pre-processing method to existing TCR repertoire data derived from 32 mice.^32^ For each mouse, two samples were collected from the spleen: one from the T cell zone (TCZ) and one from the B cell zone (BCZ). Sixteen samples were obtained from control mice injected with 200μl phosphate-buffered saline (PBS) and sacrificed after 3 days. The other sixteen samples were obtained four days after immunization with 200μl PBS with 10^9^sheep red blood cells (SRBC).

### Similarity-based weighting

For the simulated and real TCR-seq data, we first created similarity matrices to adjust raw read counts based on sequence similarity of the CDR3β sequences. Let **S** denote all sequences present across all samples. We define a | **S** | *×* | **S** | matrix **M** with the similarity value between each pair of sequence *s*_*i*_ and *s*_*j*_ . We then derive neighbouring sequences by applying a threshold *t* to **M**. For each sample, let the vector **C** then be the raw counts data over **S**. Our pre-processing step updates the counts for a sequence *s*_*i*_ with a weighted sum over the counts of similar sequences (**Figure 1)** as follows:

**Figure 1.**
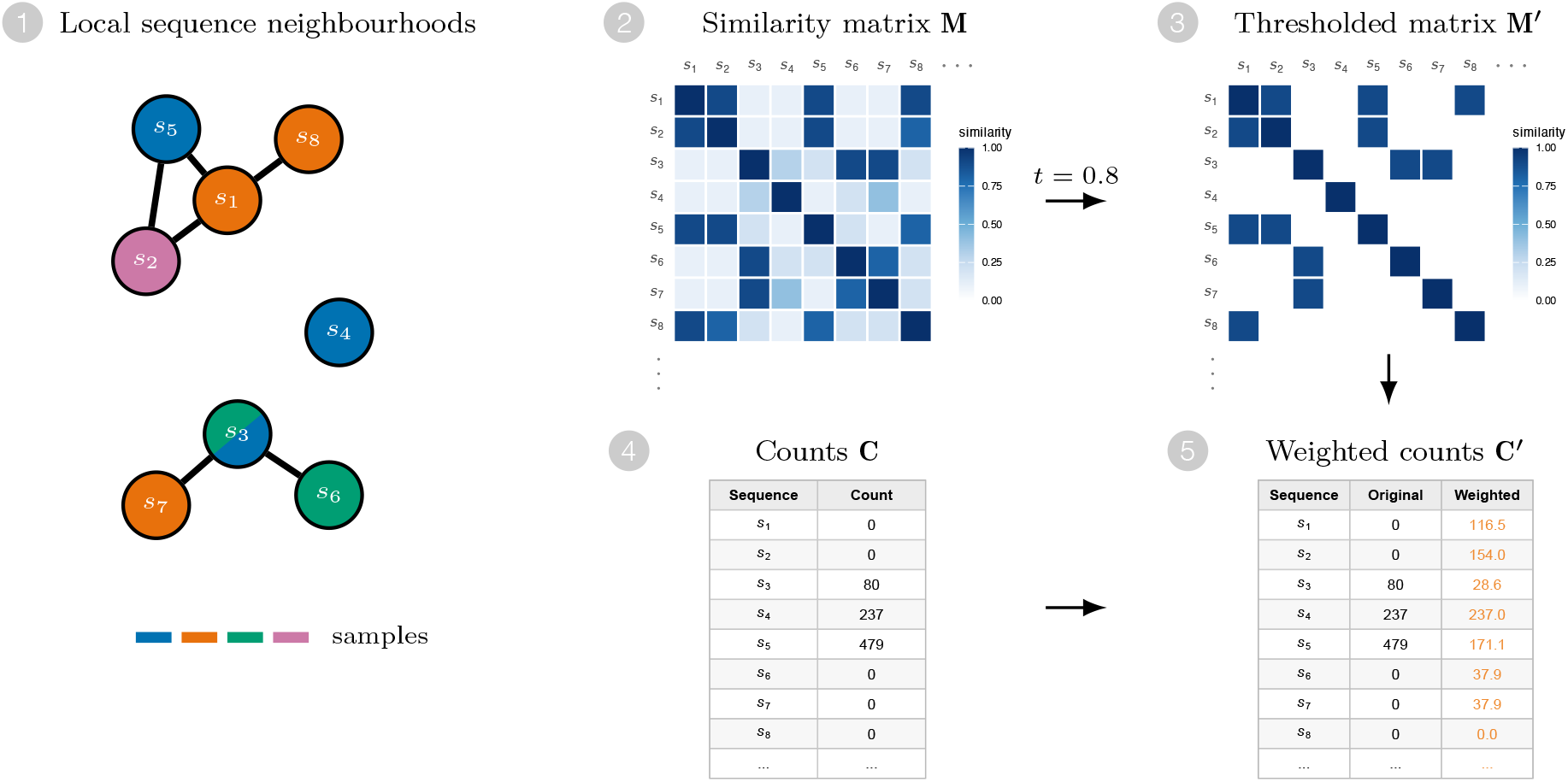
Schematic of similarity-based counts weighting. Example with four samples and eight sequences, some of which are similar. (1) Local neighbourhood of the eight sequences, coloured by sample of origin. Most sequences are private to a single sample, but *s*_3_ (split fill) is found in two samples. (2) Similarity matrix *M*, derived using the Hamming based metric. (3) Thresholded matrix *M*^*′*^, with only similarities above *t* = 0.8. (4) Raw counts vector *C* for one example sample (blue in (1)). (5) Weighted counts vector *C*^*′*^, obtained using *M*^*′*^to average counts over sequence neighbourhoods.

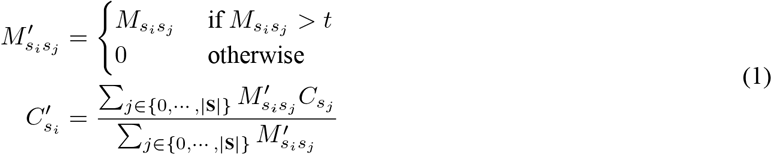

We used two methods to calculate the similarity matrix: one based on the Hamming distance between sequences and the other based on the BLOSUM62 alignment^33^ between sequences. We processed the dataset by grouping sequences according to their V and J segments and their lengths and calculating a similarity matrix for each subgroup. Earlier research has shown that high-performance clustering of epitope-specific TCR sequences can be achieved when considering only sequences of the same length.^34^ Moreover, restricting comparisons to sequences of equal length not only aligns with biological relevance but also reduces computational costs, making this approach both scalable and efficient for large TCR-seq datasets.

We define the Hamming-based similarity metric by normalizing over sequence lengths and converting the distance matrices into similarity matrices by subtracting the normalized distances from one. The Hamming similarity for the pair of sequences *s*_*i*_ and *s*_*j*_ is thus defined as 1−(*h*_*i,j*_/*n*_*char*_), with *h*_*i,j*_ the Hamming distance and *n*_*char*_ the length of the CDR3 amino acid sequence.

For the BLOSUM62 approach, we took the original target probabilities^33^ and normalized per amino acid to get the probability vector per row. Then, for each combination of amino acids, we computed the squared distance between probability vectors, and normalized to bound the score by [0,1]. The similarity score for a sequence is then defined as the sum of distance scores normalized by the length and subtracted from one, similar to the Hamming similarity score.

Both metrics thus range from 0, indicating no similarity, to 1, indicating identical sequences. In the Hamming-based method, sequences with a similarity score above 0.8 were considered highly similar, while in the BLOSUM62-based method, sequences with a normalized alignment score above 0.9 were considered similar. For the Hamming distance method, a similarity score above 0.8 indicates that at least 80% of the sequence positions are identical. This value takes into account minor variations relative to the length of the sequence. For the BLOSUM62 method, a slightly lower threshold of 0.9 was chosen empirically.

We made our similarity-based weighting method available in the R package simweightR (https://github.com/thomcsmits/simweightR).

### Differential abundance analysis

To study how neighbouring information influences DA analysis on TCR-seq data, we compared the results of three methods: two parametric, DESeq2, edgeR, and one non-parametric, the Wilcoxon test.

The count matrix of sequence read counts per amino acid junction sequence, V gene and J gene combination, as well as a condition label vector, were used as input to all methods. Sequences occurring in only one mouse were removed before applying the similarity-based weighting since those are less likely to reflect biologically meaningful patterns across conditions.

For edgeR (v4.0.9), normalization using the trimmed mean of M-values (TMM) was applied. Then, a quasi-likelihood generalized linear model was fitted, followed by a quasi-likelihood F-test to identify the differential clones. In DESeq2 (v1.42.0), normalization, dispersion estimation, and DA analysis were performed using the *DESeq* function included in the package. Since the Wilcoxon test is not a regression-based method, it cannot adjust for confounding factors, such as sequencing depth. Therefore, the matrix was normalized before applying the Wilcoxon test using counts per million (CPM) via edgeR’s *calcNormFactors* function and transformed into a normalized count matrix.^9^ Then, a Wilcoxon rank-sum test was conducted for sequence to compare its expression levels between treatment and non-treatment samples.

### Statistical analysis

In the simulated TCR-seq datasets, the true positives were known and thus the TRP and FPR can be calculated. We used the simulated datasets to test the calibration of the tests. With a p-value cut-off of 5%, we expect the FPR to be 5%, and can thus test this calibration.

For the real TCR-seq datasets, true positives are not known. We can, however, compare the number of found differential clonotypes with an estimation of the false positives. In the differential abundance testing, p-value correction was applied with the Benjamini-Hochberg method.^6^ Since averaging counts within samples based on their neighbourhood makes instances dependent on each other, not all p-value correction methods are applicable. The Benjamini-Hochberg method can be used for dependent samples if positively correlated. Here, we assume a positive correlation as we adjust the read count based on the premise that sequences that have a higher sequence similarity are more likely to bind the same epitope.

To estimate the false positives, we used a label-shuffling approach. Half of the sample labels (naïve or SRBC) from each group were randomly reassigned to the opposite group to break any true biological association between TCR expression and condition labels. DA analysis was performed on these shuffled datasets, and the resulting differential clonotypes were used to estimate the FPRs.

All statistical analyses were performed within the R platform for statistical computing version 4.3.1.^35^ 6

## Results

### T cell receptor sequencing data has different characteristics compared to other RNA sequencing data

Since DGE methods are designed for RNA-seq, we wanted to investigate how bulk TCR-seq data differs from bulk RNA seq data by comparing the proportion of zero values across bulk RNA-seq, single-cell TCR-seq (scTCR-seq), and bulk TCR-seq datasets (**Figure 2A**). Bulk RNA-seq, which captures the average gene expression across a population of cells, exhibited relatively few zeros, with an average of 35.92% zeros per sample *±* 3.5 (n=109).^36^ In contrast, scTCR-seq, which measures expression at the single-cell level, showed a much higher prevalence of zeros, with an average of 83.20% zeros *±* 5.02 (n=2).^37^ This sparsity arises due to biological heterogeneity–most clonotypes are private and not share dbetween individuals–as well as technical factors such as low sequencing depth. Bulk TCR-seq data exhibited the highest proportion of zeros, with an average of 96.76% zeros *±* 2.23 (n=66). This extreme sparsity primarily reflects the absence of certain TCR clonotypes within a sample but can also be attributed to sampling or sequencing limitations. Methods developed for analyzing RNA-seq datasets might, therefore, not fit scRNA-seq and TCR-seq data since data assumptions are not met.

**Figure 2.**
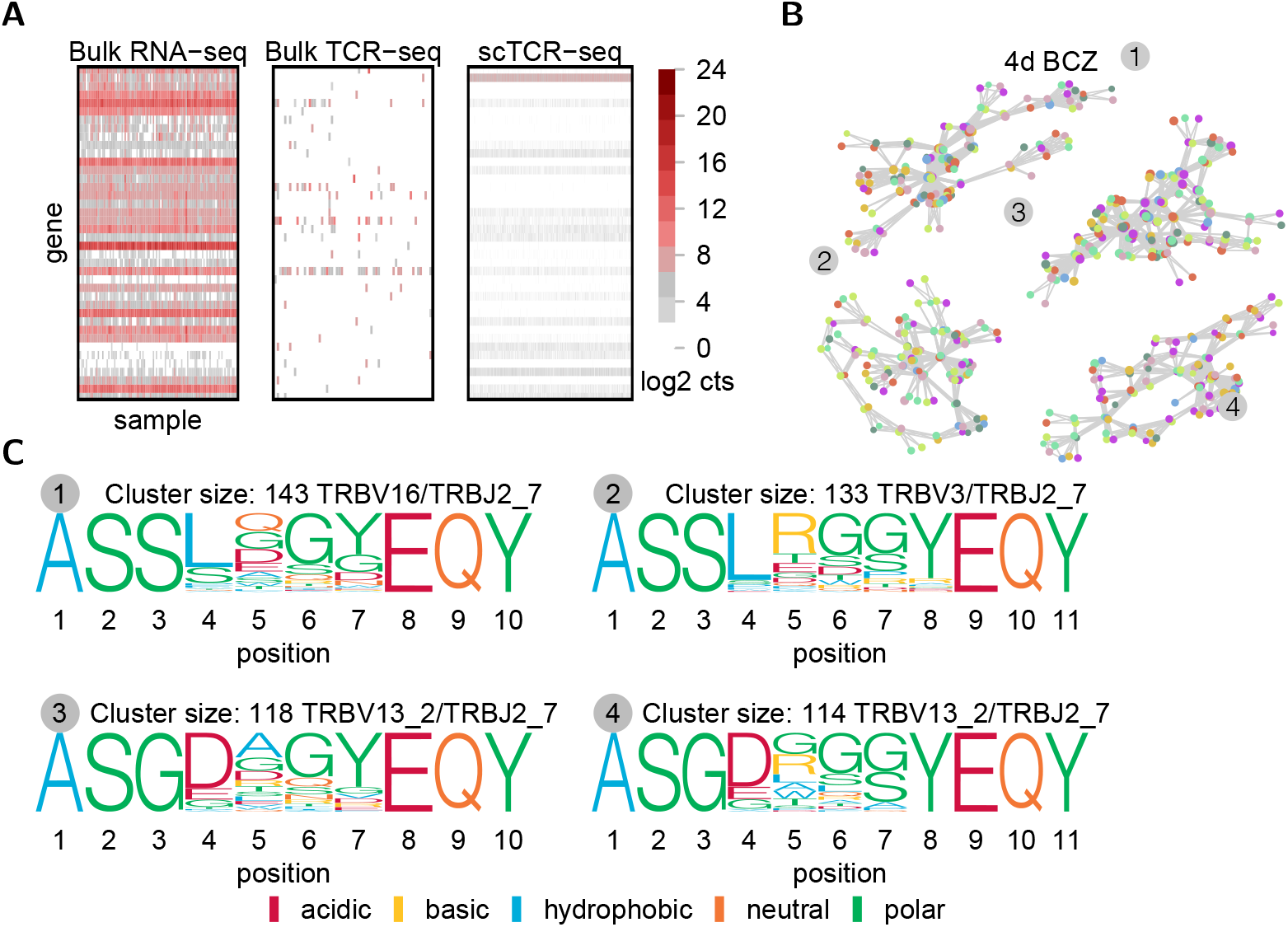
While TCR sequencing data is sparse, it contains clusters of similar sequences. **(A)** Comparison of T cell receptor sequencing data,^32^ RNA-bulk sequencing data,^36^ and single-cell sequencing data.^37^ From each dataset, the first 40 entries are visualized. **(B)** The four largest sequence clusters in TCR repertoires from 8 mice, 4 days post-infection with SRBC from the B cell zone (BCZ). Sequences are considered neighbours if their Hamming distance is no greater than 1 and they share the same V gene and J gene. Each dot represents a TCR sequence, with the size corresponding to the read count. Different colors represent individual mice. **(C)** Sequence logos of the clusters shown in panel B, where the height of each position reflects the probability of an amino acid occurring at that position.

Despite its sparsity, TCR-seq data reveals structured patterns, with TCR sequences forming clusters based on sequence similarity (**Figure 2B, C**). In TCR repertoires from mice infected with SRBC, we identified several large clusters of closely related TCR sequences across different mice. These clusters represent groups of clonotypes that share the same V and J gene segments and differ by only a single amino acid. Such clustering suggests that clonotypes with high sequence similarity are functionally related and may arise due to convergent recombination or selection by the immune system.^23, 25^ However, due to the data’s sparsity, most sequences within these clusters appear in only one mouse, complicating DA analysis. Nevertheless, since similar sequences are likely to bind the same epitope, we could leverage information from similar sequences in other mice to mitigate data sparsity.

### Local weighting improves detection of differentially abundant TCRs in simulated repertoires

To investigate the impact of sparsity on DA analysis and assess whether incorporating information from neighbouring instances influences the true positive rate (TPR) and false positive rate (FPR), we first used simlated data. We generated 90 simulated repertoires using immuneSIM^29^ and added known reactive sequences from the VDJ database^30^ to 45 of those (**Figure 3A**). These sequences were modeled for proliferation using a Galton-Watson process, in which each sequence produced two offspring with a probability of 0.3. This process iterated until no additional offspring were generated, simulating clonal expansion of reactive sequences (**Figure 3B**).

**Figure 3.**
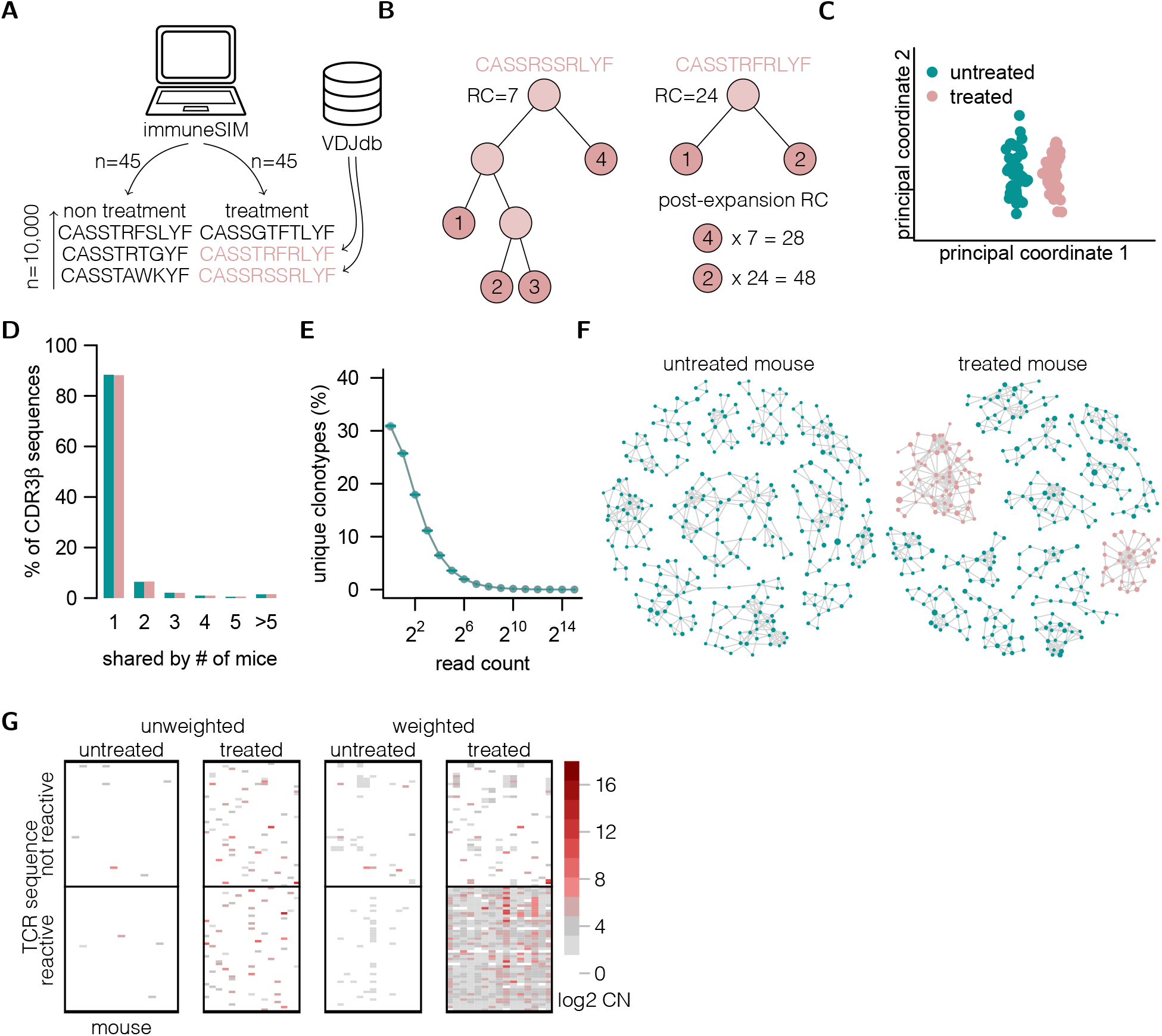
Simulated T cell receptor CDR3β data, enhanced with reactive sequences, reveals distinct clusters of similar CDR3β sequences. **(A)** We generated 90 immune repertoires using immuneSIM, incorporating sequences from the VDJ database into 75 of them to represent the treatment group. **(B)** The proliferation of cells in the reactive sequences was modeled using a Galton-Watson process. In each evolutionary step, each cell produced two offspring with probability *p*. The simulation continued until no new offsprings were generated in an evolutionary step, marking the termination of the process (RC: read count). **(C)** Multidimensional scaling overview of the simulated repertoires, based on overlap (Jaccard index) in junction aa sequence, V region, and J region. **(D)** Sharing distribution of CDR3β sequences in the simulated repertoires, based on the junction aa sequence, V region and J region. **(E)** Read count distribution of the treated and untreated samples. **(F)** Clustering plots illustrating the simulated data from one untreated and one treated repertoire, where each dot represents a unique TCR sequence. Connections between dots indicate a maximum Hamming distance of 1, and dot sizes reflect read counts. **(G)** Read counts of 100 reactive sequences in 15 treated and 15 untreated samples. Left: unweighted read counts. Right: locally weighted read counts, considering sequences with close Hamming distances in the neighbourhood. Color scales are logarithmic, adding 1 to each read count. In each panel the first 100 sequences in the dataset are shown.

To assess repertoire similarity, we calculated the Jaccard index based on shared amino acid junction sequences, V genes, and J genes. Multidimensional Scaling (MDS) of the data revealed distinct clustering of repertoires by treatment status, indicating increased similarity among treated samples due to reactive sequence enrichment (**Figure 3C**). However, since only a few sequences were reactive, these changes did not drastically alter the overall repertoire composition. This aligns with the goal of our simulation, where the treatment-induced enrichment of reactive sequences did not significantly impact the broader diversity of the TCR repertoires. Furthermore, the treated samples showed no substantial increase in the overall abundance distribution, as depicted in **Figure 3E**, which is consistent with the limited effect of reactive sequence proliferation on the overall repertoire.

After generating TCR-seq data, we applied local weighting. We clustered the sequences in one treated and one untreated repertoire (**Figure 3F**), where each cluster represented TCR sequences with minimal sequence divergence (Hamming distance ≤ 1), and larger dot sizes indicated higher read counts. In the treated samples, reactive sequences formed distinct clusters that included both low- and high-abundance sequences, confirming the effectiveness of the proliferation model. As a result, weighting the read counts based on these local neighbourhoods produced a more even distribution of read counts for the reactive sequences in the treatment group (**Figure 3G**).

Applying local weighting based on Hamming distance and BLOSUM62 alignment in all three methods improved the detection performance of differential expression methods. TPRs increased for edgeR, DESeq2, and the Wilcoxon test (**Figure 4A**). The influence of local weighting on the FPRs varied between the methods as shown in **Figure 4B**. For edgeR, FPR increased to approximately 15% with Hamming-based weighting and 27% with BLOSUM62-based weight ing. DESeq2 maintained FPRs below its claimed threshold of 5%, with slight increases after weighting. The Wilcoxon test showed low FPRs for the unweighted sequences, which increased to 5% after weighting, in line with its claimed FPR threshold. Comparing the log2 fold changes (log2FC) before and after applying local weighting shows that the weighting process enhanced the fold changes for reactive sequences (shown in pink), as most of them shifted to the left of the diagonal line. The adjustment amplified signal detection while reducing noise, further improving the differentiation between treated and untreated groups (**Figure 4C**).

**Figure 4.**
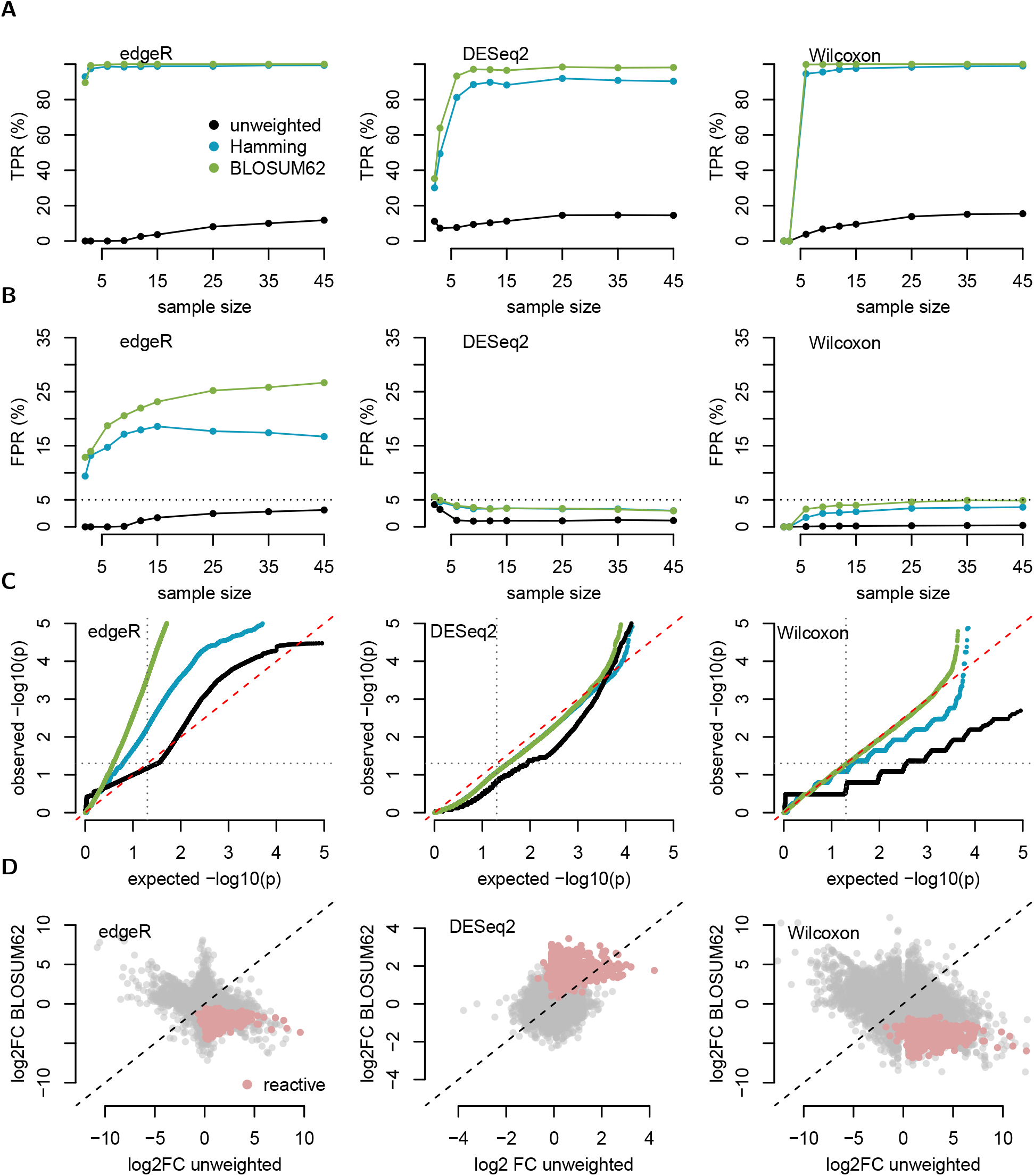
Local weighting on simulated T cell receptor data improves the true positive rate. **(A)** True positive rate of edgeR, DESeq2, and the Wilcoxon test for varying sample sizes calculated on the simulated dataset described in Figure 3, with local weighting based on Hamming distance and BLOSUM62 alignment. **(B)** False positive rate of edgeR, DESeq2, and the Wilcoxon test for different sample sizes and local weighting on the simulated dataset described in Figure 3. **(C)** Quantile-quantile plot showing expected versus observed p values.**(C)** Fold changes in sequences, before and after applying local weighting using the Hamming neighbourhood for a sample size of 15. From the unreactive sequences we randomly sampled 20,000 points to plot.

Our findings demonstrate that local similarity-weighting strategies improve the detection of differentially abundant TCRs in simulated data. The Wilcoxon test, in particular, maintained more reliable FPR control compared to parametric methods, which were less stable with and without local weighting applied.

### Local weighting as a preprocessing step improves power and calibration of differential TCR expression analyses in real data

Building on the performance evaluation of our methods using simulated TCR-seq data, we next applied the methodology to a real TCR-seq dataset. This dataset includes 32 mice divided into two groups: 16 naïve (8 samples each from the BCZ and TCZ) and 16 sampled four days post-infection with SRBC (8 BCZ, 8 TCZ). Using an MDS plot based on the Jaccard index, we visualized the overlap between samples (**Figure 5A**). The increased heterogeneity of real TCR-seq datasets compared to simulated data is evident, with samples showing substantial overlap between naïve and SRBC-infected groups, reflecting the inherent complexity introduced by biological variability, sequencing depth differences, and sampling biases. This overlap complicates DA analysis.

**Figure 5.**
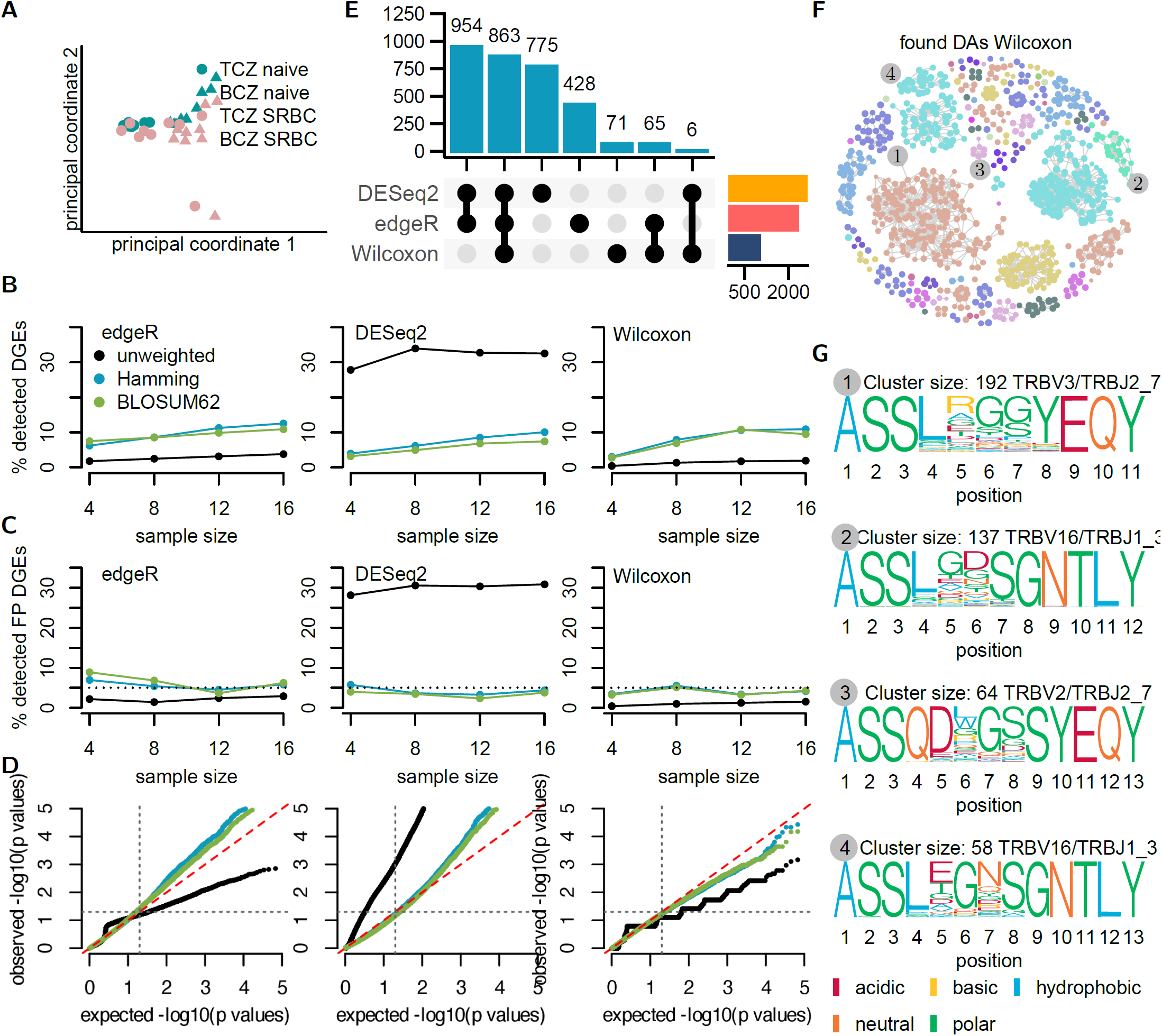
In real TCR-seq data, the Wilcoxon test remains stable while weighting improves detection and calibration. **(A)** Multidimensional Scaling overview plot of CDR3β sequences, based on overlap (Jaccard index) in junction aa sequence, V region, and J region. The data were obtained from 32 mice, 16 naïve (8 BCZ, 8 TCZ) and 16 4d after infection with SRBC (8 BCZ, 8 TCZ). **(B)** Percentage of detected differentially abundant clonotypes found with edgeR, DESeq2, and the Wilcoxon test shown for varying sample sizes (based on the number of mice) and without weighting, as well as with Hamming-based and BLOSUM62-based similarity weighting. **(C)** Percentage of detected differentially abundant clonotypes found with edgeR, DESeq2, and the Wilcoxon test after shuffling the sample labels. **(D)** Quantile-quantile plot showing the expected versus observed p values (after Benjamini-Hochberg correction) for edgeR, DESeq2, and the Wilcoxon test on the data with shuffled labels. **(E)** Intersections of differentially abundant TCRs identified by edgeR, DESeq2, and the Wilcoxon test after p value correction. **(F)** Differentially abundant TCRs identified using the Wilcoxon test with Hamming-based similarity weighting, where sequences are connected if their Hamming distance is no more than 1. **(G)** Sequence motifs of the four largest clusters of differentially abundant TCRs from F. The height of each amino acid is proportional to its occurrence probability at that position in the cluster.

Next, we examined the number of detected differential clonotypes using the Wilcoxon test, edgeR, and DESeq2 with and without local weighting for different sample sizes. The number of detected differential clonotypes increased with similarity weighting for the Wilcoxon test and edgeR, which showed improved detection after Hamming-based and BLOSUM62-based similarity weighting (**Figure 5B**). In contrast, DESeq2 exhibited a decrease in the number of detected differential clonotypes after applying weighting, which may indicate a reduction in false positives but also a potential loss in detection power.

To evaluate robustness, we randomly swapped half of the sample labels between groups, ensuring that the biological meaning was removed from the data. We then re-ran the analysis to estimate the FPRs of the different methods. The results for DESeq2 and edgeR on real data differed from those observed with simulated data (**Figure 4B**), indicating their sensitivity to data distribution. DESeq2 showed a high FPR of about 30%. However, when local weighting was applied, the FPR for DESeq2 decreased to around 5% for both the BLOSUM62-based and Hamming-based weighting. Contrary to the findings from the simulated data, edgeR demonstrated a relatively low FPR without weighting, which with weighting also stabilized around 5%. This indicates that the performance of both edgeR and DESeq2 can vary considerably depending on the specific dataset used.

In contrast, the Wilcoxon test showed consistent FPR patterns between real and simulated data. It maintained low FPRs for unweighted sequences, which increased to 5% after weighting. Quantile-quantile (QQ) plots further illustrated differences in calibration across methods (**Figure 5D**). For the Wilcoxon test with similarity weighting, observed p values closely matched the expected distribution, suggesting proper calibration. In contrast, both edgeR and DESeq2 displayed systematic deviations, with an excess of small p-values, indicating a potential overestimation of significant differential clonotypes.

Using a sample size of 16, the Wilcoxon test identified 1,005 differential clonotypes, while edgeR found 2,310 and DE Seq2 found 2,598 after applying Hamming-based weighting and p value correction (false discovery rate [FDR] < 0.05; **Figure 5E**). DESeq2 and edgeR showed a high overlap in their findings and detected more sequences than the Wilcoxon test. Of the differentially abundant sequences identified by the Wilcoxon test, only 71 were exclusive to this method. In contrast, edgeR identified 428 sequences, and DESeq2 detected 775 sequences that were not found by the other methods.

Finally, we analyzed clusters of differential TCRs detected by the Wilcoxon test with Hamming-based similarity weighting. Sequences were connected in a network if their Hamming distance was ≤ 1, revealing distinct clusters of related TCRs (**Figure 5F**). Sequence motifs from the four largest clusters showed strong positional conservation, with amino acid heights proportional to their occurrence probabilities (**Figure 5G**). These motifs provide insights into the functional specificity of the differential TCRs.

In summary, our results demonstrate that incorporating local similarity weighting with the Wilcoxon test improves the detection and calibration of differential TCRs in real datasets. Compared to traditional methods like edgeR and DESeq2, this approach improves robustness and uncovers biologically meaningful patterns, such as sequence motifs and clusters in TCR-seq data.

## Discussion

Traditional DGE methods, such as DESeq2^8^and edgeR,^7^are powerful tools for analyzing RNA-seq data. However, they encounter challenges when used in DA analysis on TCR-seq data, primarily due to the sparsity of TCR-seq data, with many sequences exhibiting low counts or even zeros due to both technical and biological factors.^20, 38^ These characteristics make it challenging for traditional methods, which operate under strict assumptions about the underlying data distribution, to detect differentially abundant TCRs accurately. Here, we show that these are not well-calibrated for DA analysis on TCR-seq data. Even non-parametric approaches, such as the Wilcoxon test, face challenges in TCR-seq datasets due to the abundance of zeros, which results in a high number of ties in rank calculations. Earlier methods such as ALICE^27^ and TCRnet^39^ have been developed to identify clusters of epitope-specific sequences by comparing them to a baseline repertoire. These approaches use sequence similarity to detect antigen-reactive TCR clusters. However, one challenge with these methods is their limited sensitivity to low-frequency sequences, leading to missing rare but potentially important TCRs. This limitation underscores the need for improved strategies that can effectively account for the specific characteristics of TCR-seq data.

To address these issues with TCR-seq dataset analysis, in this study, we proposed a pre-processing step for TCR-seq data utilizing local weighing methods to enhance DA analysis for TCR-seq data. Incorporating local weighting methods allowed us to address the specific challenges.

A similar approach is proposed by Yohannes et al.^40^ which constructs clusters of TCR sequences and tests for differential abundance of the clusters. However, it is more computationally expensive, and our method is more flexible for different DA analyses.

Our comparison of DA methods highlighted the differing performance of traditional parametric and non-parametric ap proaches. The Wilcoxon test performed best. It showed consistent performance across both simulated and real datasets. Unlike parametric methods, the Wilcoxon test is robust against non-normal distributions and outliers, making it suitable for sparse and skewed datasets. Incorporating local weighting helped address problems with ties in rank-based compar isons by integrating relevant biological similarity information. As a result, there was an improvement in calibration and sensitivity.

For the parametric methods, we also observed improved detection and calibration when using the weighted preprocessing in combination with DESeq2. Specifically in the mouse dataset, before weighting DESeq2 had an inflated false positive rate, which was corrected for with weighted. However, for edgeR, both unweighted and weighted counts were not consistent, indicating that edgeR may not be suitable for TCR-seq data. Parameter optimization can help to effectively address issues of sparsity and overdispersion, although this is labor-intensive and does not always generalize well across different datasets.

Both Wilcoxon and DESeq2 performed well with weighted sequence counts. However, since DESeq2 may be more sensitive to changes in data distribution, we recommend the Wilcoxon test in combination with similarity-based counts weighting for DA analysis in TCR-seq data, for it’s robustness, simplicity, and minimal need for parameter tuning.

We utilized two sequence similarity matrices to determine the local neighbourhood for weighting: one based on the Hamming distance and another based on the BLOSUM62 alignment.^33^ This approach introduces a new parameter that defines which sequences are considered part of the local neighbourhood. One limitation of our study is the absence of systematic optimization for this parameter. A potential improvement would be to conduct a grid search and evaluate performance on the simulated dataset to identify optimal thresholds. While our results show improvements with the chosen similarity thresholds (0.8 for Hamming distance and 0.9 for BLOSUM62), fine-tuning these thresholds based on dataset-specific characteristics could further enhance performance.

Another potential area for improvement is the choice of substitution matrices. While BLOSUM62 has been widely used in sequence comparison, its applicability to CDR3β sequences may not fully capture their functional nuances. Recent advancements, such as the TCR-specific substitution matrix proposed by Postovskaya et al.,^41^ could be promising. A specialized matrix could enhance our ability to detect TCRs with small sequence differences that still share epitope specificity and improve local weighting.

In conclusion, this study shows that using local weighting strategies in DA analysis can enhance the sensitivity and calibration of TCR-seq data analysis. The combination with the Wilcoxon test, in particular, offers an effective framework for controlling FPRs while utilizing the additional information obtained from local neighbourhood similarities. Our findings provide a foundation for creating more accurate and reliable TCR-seq analysis tools. Future research should focus on optimizing the parameters for local weighting, exploring alternative substitution matrices specifically designed for TCR sequences, and integrating these methods into frameworks for biological interpretation. Additionally, it would be interesting to see how this approach performs on other sparse datasets where correlations between entries are known to hold, such as single cell data.^42^ These advancements will further reveal insights into the immune system’s response to pathogens, vaccines, and autoimmune diseases, ultimately deepening our understanding of adaptive immunity.

## Acknowledgements

The authors thank Inge Wortel for valuable feedback on the manuscript. This research was supported by an NWO Vidi grant (VI.Vidi.192.084) to JT. We thank Radu-Florian Diaconescu for work on the intial version of the R package.

## Author contributions

JT and FB conceived the project. TCS and FB performed the experiments and analyzed the data with help from AA under supervision of JT. TCS created the R package. All authors discussed the results and contributed to the final manuscript.

## Data and code availability

The source code and data used to reproduce the results and analyses presented in this manuscript are available on Zenodo.^43^ The simweightR package is available at https://github.com/thomcsmits/simweightR.

